# The murine lung microbiome is dynamic and transient

**DOI:** 10.64898/2026.05.18.725910

**Authors:** Liubov Nikitashina, Kerren Volkmar, Maria Straßburger, Sascha Schäuble, Zoltán Cseresnyés, Eric Unger, Ilse D. Jacobsen, Marc Thilo Figge, Gianni Panagiotou, Thorsten Heinekamp, Axel A. Brakhage

**Affiliations:** Department of Molecular and Applied Microbiology, Leibniz Institute for Natural Product Research and Infection Biology (Leibniz-HKI), 07745 Jena, Germany; Institute of Microbiology, Friedrich Schiller University, 07743 Jena, Germany; Transfer Group Anti-infectives, Leibniz-HKI, 07745 Jena, Germany; Department of Microbiome Dynamics, Leibniz-HKI, 07745, Jena, Germany; Department of Applied Systems Biology, Leibniz-HKI, 07745 Jena, Germany; Department of Microbial Immunology, Leibniz-HKI, 07745 Jena, Germany; Cluster of Excellence Balance of the Microverse, Friedrich Schiller University, 07743 Jena, Germany

**Keywords:** Lung microbiome, commensal lung bacteria, host-mediated colonization control, phagocytosis, *Ligilactobacillus murinus*, fluorescent labeling, *Aspergillus fumigatus*, precision-cut lung slices, mice

## Abstract

**Background:** Whether the lung microbiome represents a stable microbial colonization or a transient ecosystem shaped by continuous microbial turnover and controlled by host immunity remains unresolved. The murine lung microbiome largely consists of species from the former *Lactobacillus* genus with *Ligilactobacillus murinus* as a dominant species, bacterial genera such as *Streptococcus*, *Staphylococcus*, *Mammaliicoccus*, *Enterococcus* and other less frequently detected bacteria. Here, we directly addressed the question of persistence and host interaction of a dominant murine lung commensal *in vivo* and focused on the host immune response towards lung commensal bacteria.

**Results:** We developed a transformation strategy for stable genomic integration of a green fluorescent protein (GFP)-encoding gene to track the fate of a lung bacterium. Following intranasal administration of GFP-labeled *L. murinus* in mice, bacteria were readily detected in the lungs at early time points but declined rapidly and became undetectable after 72 hours, as determined by quantification of viable bacteria and qPCR. Flow cytometry and fluorescence imaging revealed efficient uptake of GFP-labeled bacteria by lung phagocytes. These findings indicate that even dominant members of the murine pulmonary microbiota normally detected at low abundances are transiently present in the lungs without causing infection. We further analyzed the effects of moderate and high bacterial concentrations. While moderate bacterial loads were efficiently controlled without clinical effects, high concentrations induced severe lethargy, indicating a threshold-dependent host response. Finally, we demonstrated that pulmonary commensals such as *L. murinus*, *Staphylococcus xylosus*, and *Mammaliicoccus sciuri*, as well as conidia of the opportunistic lung pathogen *Aspergillus fumigatus*, are phagocytosed at comparable rates in macrophage assays.

**Conclusions:** Our data demonstrate that even lung-adapted bacterial species fail to establish stable colonization and are instead subject to rapid immune-mediated elimination contributing to the maintenance of a low microbial burden in the lungs. While this homeostatic balance supports health, elevated bacterial loads trigger immune activation and, at high levels, lead to health deterioration. Together, these results support a model of a highly dynamic and transient lung microbiome, maintained by continual microbial immigration rather than long-term colonization. Accounting for the lung microbiome dynamics is essential for understanding host-microbiota interactions and respiratory health.

## Background

The lungs are responsible for the vital function of oxygen exchange. Due to their function, they remain continuously exposed to microorganisms from the air through breathing as well as from the upper airways. To maintain homeostasis, acquired microorganisms are cleared through mechanical defenses as well as innate and adaptive immune responses. Disruption of the lung homeostasis can lead to disorders, with chronic obstructive pulmonary disease (COPD) and lower respiratory tract infections representing the leading causes of mortality worldwide [1, 2]. Despite effective clearance mechanism, healthy lungs have been shown to harbor a distinct microbial community contrary to the long-held view of sterility [3, 4]. Increasing evidence indicates that the lung microbiome plays a critical role in maintaining pulmonary homeostasis, and its dysregulation is associated with chronic lung diseases, including COPD, as well as acute infections such as SARS-CoV-2, pneumonia, and invasive aspergillosis [5–15]. Therefore, in certain cases, the lung microbiome can serve as an indicator of disease and even as a diagnostic marker [16]. A few studies have examined the functional contributions of individual members of the lung microbiome, demonstrating their capacity to modulate host immune responses, through both pro- and anti-inflammatory mechanisms [17–23]. However, our understanding of how the host regulates the lung microbiome in health and disease remains limited.

The composition of the lung microbiome has been shown to resemble the composition of the oral microbiome [24, 25] and is characterized by a very low overall microbial burden and high interindividual variation in community composition [24, 26]. These characteristics, in combination with the physiological features of the respiratory tract and limited nutrient availability [27], led to the hypothesis that the formation of the lung microbiome community follows an “adapted island model” [25, 26]. According to this model, the lung microbiome is primarily shaped by the acquisition of microorganisms from the oral cavity and oropharynx *via* microaspiration and their subsequent elimination through mucociliary clearance as well as innate and adaptive immune mechanisms [25, 26, 28]. In turn, the local microbial proliferation in the lung is suggested to have a minimal impact on the community structure formation [25, 26, 29]. Although, this hypothesis was consistent with microbial metabolic activity analysis [30], the experimental proof remained lacking. In turn, several studies suggested the existence of a relatively stable core lung microbiome [9, 31]. Thus, longitudinal analyses based on sequential bronchoscopy demonstrated that within-individual microbial composition remained more stable over time than would have been expected by stochastic uptake, suggesting a persistent bacterial subset [31]. Yet, it remains unclear whether members of the lung microbiome can persist and propagate in the lung environment. They may also possess mechanisms that reduce recognition by phagocytes leading to a less efficient clearance from the lungs.

We previously observed that the lung microbiome of specific pathogen-free (SPF) laboratory mice was largely dominated by *Ligilactobacillus murinus* [32], which was in line with several other studies [23, 33, 34]. One of the possible mechanisms responsible for high prevalence of *L. murinus* in murine lungs might be its adjustment to the lung environment and ability to persistently colonize lungs.

Based on our previous findings, in this work, we address the fundamental question whether bacteria detected in the lung microbiome can persist in the lungs and form a resident community or whether the lung microbiome rather represents a transient bacterial community tightly controlled by the host. For this, we colonized the lungs of specific-pathogen-free (SPF) mice with *L. murinus*, the dominant and representative member of the lung microbiome. In order to distinguish the intranasally administered bacteria from bacteria already present in the lung microbiome, for colonization, *L. murinus* was genetically modified to stably produce the green fluorescent protein (GFP). This approach enabled us to track colonization over time, localize the bacteria within the lungs and analyze their internalization by host lung cells.

## Methods

### Experimental animals

Specific-pathogen-free (SPF) female BALB/c mice (7 to 10 weeks old) were obtained from Janvier Labs, France. Mice were housed in groups of 5 animals in individually ventilated cages and fed with normal mouse chow and water *ad libitum*. All animals were handled in compliance with the European animal welfare regulations. Approval for the experiments was obtained from the responsible federal/state authority and the ethics committee, which are based on the provisions of the German Animal Welfare Act (permit no HKI-24-003).

### Microorganisms and culture conditions

*Ligilactobacillus murinus* MAM1902, *Staphylococcus xylosus* MAM1901, and *Mammaliicoccus sciuri* MAM2035 were previously isolated by us from the lower respiratory tract of C57BL/6 mice [32]. *L. murinus* was cultured on Columbia Blood Agar (CBA) (Sigma-Aldrich, Germany) when required supplemented with 0.8 µg/ml or 2 µg/ ml of erythromycin (Er), or with 6 µg/ml or 8 µg/ml chloramphenicol (Cm) at 37 °C aerobically or anaerobically and in Man-Rogosa-Sharpe (MRS) medium (Carl Roth, Germany) at 37 °C and 75 rpm aerobically. *S. xylosus* and *M. sciuri* were cultured on Tryptic Soy Agar (TSA) at 37 °C aerobically. *A. fumigatus* CEA10 [35] (wild type) was cultured on *Aspergillus* minimal medium [36] (AMM) agar at 37 °C aerobically. NEB Turbo Competent *E. coli* were cultured in LB or on LB agar supplemented with 200 µg/ml Er.

### Bacteria whole genome sequencing and annotation

#### Isolation of genomic DNA

Whole genome sequencing and annotation were performed for *L. murinus* MAM1902. Lysis of bacteria was performed with InnuPREP Bacteria Lysis Booster (Analytik Jena, Germany) according to the manufacturer’s protocol. Isolation of gDNA was performed with the NucleoBond HMW DNA kit (Macherey-Nagel, Germany) according to the protocol for ‘Enzymatic lysis’. gDNA purification was finalized using the NucleoSnap Finisher Midi Kit (Macherey-Nagel, Germany) according to the manufacturer’s protocol. DNA was eluted with 200 µl low TE buffer (New England BioLabs, USA).

#### Library construction and genomic DNA sequencing

Genomic DNA sequencing was carried out by Novogene (Cambridge, UK) using PacBio Single-Molecule Real-Time (SMRT) sequencing as follows: for the SMRTbell (Pacific Biosciences, USA) library generation, gDNA was fragmented, DNA fragments were damage-repaired, end-repaired, and A-tailed. The SMRTbell library was produced by ligating universal hairpin adapters to double-stranded DNA fragments. After exonuclease digestion and the AMPure PB beads (Pacific Biosciences, USA) purification step, sequencing primers were annealed to the SMRTbell templates, followed by binding of the DNA polymerase (Pacific Biosciences, USA) to the annealed templates and sequenced on PacBio Sequel II/IIe systems (Pacific Biosciences, USA).

#### Genome assembly, annotation, and phylogenetic analysis

PacBio raw reads were assembled using Flye v2.9.0 [37]. The assembled genomes were polished with Arrow v2.3.3 from the Genomic Consensus package (Pacific Biosciences, USA), circularized with Circlator v1.5.5 [38], polished with Arrow v2.3.3, and the assembly was assessed with BUSCO v4.0.2 [39].

Genome annotation was performed with the NCBI Prokaryotic Genome Annotation pipeline. Phylogenetic analysis was done with AutoMLST2 [40] in a Denovo mode and with standard settings.

### Generation of a GFP-producing *L. murinus* transformant strain

#### Generation of a plasmid for integration into the chromosome

The genome integration cassette consisted of the 5’ flanking region, 1000 bp of the *L. murinus* MAM1902 enolase promoter region, the gene encoding enhanced GFP derived from plasmid pTRKH3-ermGFP (addgene plasmid # 27169 [41]), the chloramphenicol acetyltransferase gene from plasmid pC194 [42], and the 3’-flanking region. 5’- and 3’-flanking regions were 1,297 and 1,245 bp long respectively and located in the inactive prophage region of *L. murinus* MAM1902 genome. The DNA fragment containing enolase promoter region, *GFP*, chloramphenicol resistance cassette, and the 3’-flanking region (4715 bp) was synthesized by Genewiz (Azenta Life Sciences, Germany) and amplified using primers Lm_Peno_F and Lm_flank_2_R. The 5’-flank region was amplified using primers Lm_flank_1_F and Lm_flank_1_R. The two DNA fragments were assembled using the NEBuilder HiFi DNA Assembly Kit (New England BioLabs, Germany) and amplified using primers pTRKH3_flank_1_F and pTRKH3_flank_2_R. The pTRKH3 backbone was amplified with primers Flank_2_pTRKH3_F and pTRKH3_R and the vector was assembled by NEBuilder HiFi DNA Assembly and the resulting plasmid pTRKH3-L.m.int.enoGFP was used for transformation of NEB Turbo Competent *E. coli (*New England BioLabs, Germany). All primers are listed in Table S1.

#### Transformation of L. murinus

*L. murinus* MAM1902 was transformed with either the expression plasmid pTRKH3-ermGFP or integration plasmid pTRKH3-L.m.int.enoGFP. For transformation of *L. murinus*, 90 ml MRS medium in a 100 ml Erlenmeyer flask were inoculated with *L. murinus* harvested from CBA plates with a starting OD_600_ = 0.2. The culture was incubated at 37 °C and shaken with 75 rpm up to an OD_600_ = 1. The culture was centrifuged and washed three times with 20 ml ice-cold 0.3 M sucrose, 10% (v/v) glycerol solution in water. The pellet was finally resuspended in 1 ml 0.5 M sucrose, 10% (v/v) glycerol solution in water. 100 µl of the bacterial suspension were mixed with 1 µl plasmid DNA containing 750 ng DNA. All procedures were performed on ice. Transformation was performed by electroporation using a 1 mm gap electroporation cuvette (Biozym, Germany) with an Evaporator (Eppendorf, Germany) set to 2,500 V. Transformants were selected on CBA supplemented with 0.8 µg/ml erythromycin by anaerobic incubation for 2 days at 37 °C.

#### Analysis of GFP insertion into L. murinus genome

GFP-positive colonies of *L. murinus* transformed with pTRKH3-L.m.int.enoGFP were passaged on CBA with or without chloramphenicol and their resistance to erythromycin and chloramphenicol was monitored. Insertion was confirmed by whole genome sequencing (Eurofins Genomics, Germany) of *L. murinus enop-GFP* using Oxford Nanopore Technology (ONT) Sequencing with Bacterial Genome Sequencing Analysis Pipeline v1.0.. Raw nanopore sequencing reads were assessed for quality and filtering. Filtlong v0.2.1 [43] was used to remove short and low-quality reads from the raw nanopore sequencing data. The high-quality nanopore sequencing reads were then used for *de novo* assembly of the bacterial genome. The assembly was performed using Flye v2.9.3 [37] with parameters optimized for bacterial genomes. The resulting contigs were further polished using Medaka v1.8 (ONT research, UK) to improve base accuracy. The annotation was performed by the prediction of coding sequences, tRNAs, rRNAs, and other genomic features based on published databases such as RefSeq and UniProt. The quality of the assembled genomes was assessed using QUAST v5.2 [44], CheckM2 v1.0.1 [45], and Mash v2.3 [46]. The purity of samples was ensured with minimap2 v2.24 [47] by mapping the sequence-cleaned reads onto the assembly and employing Clair3 v1.0.4 [48] to call variations (SNPs and INDELS) within the assembled genome. To determine the inserted sequence and the location of the insertion, pTRKH3-L.m.int.enoGFP was aligned to the genome sequence of *L. murinus enop-GFP* with Genius Prime (New Zealand). Stability of *GFP*-expression was confirmed by fluorescence microscopy using a Keyence BZ-X800 microscope (Keyence, Japan) of *L. murinus enop-GFP* cultivated on CBA without antibiotics for seven generations.

### Comparison of growth of *L. murinus* wild type and *L. murinus enop-GFP*

*L. murinus* MAM1902 (wild type) and *L. murinus enop-GFP* were cultivated in 200 µl MRS in a 96-well plate sealed with PCR foil (Thermo Fisher, Germany) without shaking. After 2 h, precultures were used to inoculate fresh cultures to a starting OD_600_ of 0.15 for monocultures and a combined starting OD_600_ of 0.3 for 1:1 *L. murinus* MAM1902:*L. murinus enop-GFP* co-cultures. OD_600_ measurement and CFU determination of colony forming units (CFU) were performed at 0 h, 24 h, 48 h, and 72 h of cultivation. Experiments were performed in triplicate.

### Administration of *L. murinus enop-GFP* to mouse lungs

To analyze the dynamics of lung bacteria presence in the lungs, *L. murinus enop-GFP* was administered to BALB/cByJRj mice and the bacterial load in the lungs was monitored over time. *L. murinus enop-GFP* was cultured on CBA anaerobically for 16 h followed by cultivation in MRS medium for 50 min at 75 rpm, 37 °C and washed twice with PBS (Gibco, UK). Final bacterial suspensions were prepared in PBS. To inoculate the lungs of mice with *L. murinus-GFP*, animals were anesthetized by an intraperitoneal anesthetic combination of midazolam, fentanyl, and medetomidine. Twenty μl of the bacterial suspension were applied to the nares of the mice. Deep anesthesia ensured inhalation of the bacterial inoculum. Anesthesia was terminated by subcutaneous injection of flumazenil, naloxone, and atipamezole. To determine the optimal dose, mice were intranasally inoculated with four different doses of bacteria, *i.e.*, 10^5^, 10^6^, 10^7^, and 10^8^. Forty-eight hours post-administration, bacterial load was quantified in two mice per dose. Due to signs of severe lethargy, mice inoculated with 10^8^ bacteria were sacrificed after 6 h for animal welfare reasons. To monitor *L. murinus enop-GFP* presence in the lungs over time, 25 mice were inoculated with 10^7^ bacteria. Samples were collected 5 min (day 0), 24 h (day 1), 72 h (day 3), 7 days, and 14 days post-administration. At these time points, mice were euthanized, and lung samples were obtained under sterile conditions.

### Quantification of *L. murinus enop-GFP* in the lungs

To obtain uniform samples, dissected lungs were transferred to gentleMACS C Tubes (Miltenyi Biotec, Germany) and dissociated with the Lung Dissociation Kit (Miltenyi Biotec, Germany) in a gentleMACS Dissociator (Miltenyi Biotec, Germany). The total volume of dissociated lung sample suspension was approximately 2.8 ml, from which an aliquot was used for CFU enumeration, 500 µl were stored at -70 °C in sterile PCR-clean Eppendorf tubes for DNA extraction, and the remaining suspension was kept at 4 °C until being analyzed by flow cytometry.

#### L. murinus enop-GFP CFU quantification

Lung suspensions were serially diluted and plated on CBA. Plates were incubated anaerobically at 37 _°_C for 48 h followed by aerobic incubation for 2 h. *L. murinus enop-GFP* numbers were evaluated by counting fluorescent CFUs. For samples from days 0, 1, and 3 post-colonization, analysis was performed in triplicates, for days 7 and 14, analysis was performed in 7 replicates. Detection of fluorescent colonies was done by imaging the plates with an IVIS Spectrum imaging system (Xenogen Biosciences, USA) using Living Image Software v.4.0 (Caliper Life Sciences, USA) for samples from days 0, 1, 3, post-administration and by fluorescent microscopy of single colonies using Keyence BZ-X800 microscope (Keyence, Japan).

#### DNA extraction and quantification by qPCR

DNA was extracted with the DNeasy Blood & Tissue Kit (Qiagen, Germany) as previously described[32]. For quantification of *L. murinus* DNA, forward and reverse primers (400 nM) and fluorescent probe (200 nM) for a gene encoding a penicillin-binding protein were used (L.m.PBP_F 5’-AAGCTTGGCGCATCGTCATC-3’, L.m.PBP_R 5’-CAACCGTGCCGTTCAAACTG-3’, L.m.PBP_probe 5’-6-FAM/CAGGTGCAT/ZEN/AATCCATCAAAGGCTTAGCCGTAGAA/IBkFQ-3’). For the quantification of *L. murinus enop-GFP* DNA, forward and reverse primers (400 nM) and a fluorescent probe (100 nM) for an GFP-encoding gene were used (GFP_F 5’-ATACAACTACAACTCCCACAAC-3’, GFP_R 5’-TGGTAAAAGGACAGGGTCATC-3’, GFP_probe 5’-6-FAM/ TCAAAGCCA/ZEN/ACTTCAAGACCCGCCA/IBkFQ-3’). For the quantification of mouse DNA, forward and reverse primers (400 nM) and a fluorescent probe (100 nM) for a gene encoding beta-actin were used (b_actin_F 5’-AGCACAGCTTCTTTGCAGCTCC-3’, b_actin_R 5’-TGGTGTCCGTTCTGAGTGATCC-3’, b_actin_probe 5’-6-FAM/AGCGGGCCT/ZEN/TCGCTCTCTCGTGGCTAGTA/IBkFQ-3’). PCR was performed with initial denaturation step at 95 °C for 3 min followed by 40 cycles of denaturation at 95 °C for 15 s and annealing/extension at 58 °C for 20 s. For the *L. murinus* and *L. murinus enop-GFP* DNA quantification in samples from day 0, ∼100 ng and in the other samples, ∼ 1 µg of the lung tissue DNA was used for each reaction. DNA content was calculated based on the standard curves obtained for the mouse gDNA, *L. murinus* gDNA, and *L. murinus enop-GFP* gDNA.

### Internalization analysis

Lung cell suspensions were treated with Red Blood Cell Lysis Solution (Miltenyi Biotec, Germany) according to the manufacturer’s protocol. The cell number was determined with a Luna Dual fluorescence cell counter (Logos Biosystems, South Korea). To analyze internalization by total lung cell population, up to 2 x 10^6^ cells were stained with fresh concanavalin A AF647 (Invitrogen, USA) with a final concentration of 10 µg/ml. To analyze internalization by lung epithelial cells, up to 1 x 10^6^ cells were blocked with FcR Blocking Reagent Mouse for 10 min (Miltenyi Biotec, Germany) followed by incubation with antibody against mouse CD326 (EpCAM) conjugated to APC (REAfinity, Miltenyi Biotec, Germany) for 30 min.

Imaging flow cytometry was conducted with Cytek Amnis ImageStreamX MkII (Cytek Biosciences, USA) using Inspire Software (Cytek Biosciences, USA). 100,000 cells were measured for each sample. For compensation, samples processed in the same way as experimental samples, from non-colonized mice and *L. murinus enop-GFP* suspension were measured. Internalization analysis was performed with Ideas software (Cytek Biosciences, USA) in two steps: first, cell debris was removed from the analysis, in the second step, the resulting population was analyzed using an Internalization Wizard. The gating strategy is described in detail in Fig. S1 and S2.

### Precision cut lung slices (PCLS)

For generation of precision cut lung slices (PCLS), mice were euthanized by ketamine/xylazine injection into the peritoneal cavity. The lung was perfused with 3% (w/v) low-melting agarose and explanted. The individual lung lobes were embedded in 3% (w/v) agarose and cut with a Compresstome (Precisionary, USA). 200 µm thick slices were transferred to 24-well plates with pre-warmed PCLS medium (advanced DMEM/F12 (Gibco, UK), 0.1% (v/v) FCS, 15 mM HEPES, 1x Glutamax (Gibco, UK), 1x penicillin/streptomycin (Gibco, UK)). For staining, PCLS were washed once with PBS and subsequently fixed and permeabilized with fix/perm buffer (PBS, 2% (v/v) PFA, 0.3% (v/v) triton-x) for 30 min at room temperature. After washing twice with IF buffer (PBS, 2% (v/v) FCS, 0.3% (v/v) triton-x), cytoskeleton and DNA were stained with Hoechst (Thermo Fisher, Germany) and phalloidin-Alexa Fluor 647 (Thermo Fisher, Germany) in IF buffer for 30 min, at room temperature in the dark. Stained PCLS were washed trice with IF buffer and stored in PBS, protected from light, at 4 °C until analysis. Immediately before image acquisition, the PBS was discarded and PCLS were overlayed with 4 drops of slowfade glass mounting medium (Thermo Fisher, Germany).

### Image acquisition and analysis

Images and Z-stacks of spots with bacteria were acquired with a Zeiss LSM 780 confocal microscope (Carl Zeiss, Germany). Images were processed with the Zeiss ZEN software. Images of Z-stacks were analyzed using the Fiji software. Channels were split, smoothed using a Gaussian blur filter and re-scaled to contain 0.35% saturated pixels. Afterwards, RGB images were created by merging the processed channels. For 2D visualizations, maximum projections were created using the “Z projection” (projection type = max intensity). For 3D visualizations, RGB images were processed using the “volume viewer” plugin, with mode set to “volume”. Snapshots of regions of interest (ROIs) were obtained with varying angles of rotation, zoom and slice position.

3D reconstructions were performed using Imaris 11.0.1 software (Bitplane, Oxford Instruments Andor, Northern Ireland). Image files were imported in their native format as z-stacks into Imaris 11.0. and converted to the Imaris IMS format. The segmentation of each channel was performed using the surface segmentation module. (I) Nuclei and bacterial DNA were identified by thresholding the DAPI channel using the Otsu method. (II) Cytoskeleton was identified by thresholding of the Alexa Fluor 647 channel. (III) EGFP-expressing bacteria were identified and distinguished from background by machine-learning-based thresholding on both the eGFP and DAPI channel. The resulting objects were filtered for size and geometry to further enhance the precision of the segmentation and then depicted as 3D surface renderings. The transparency of the cytoskeleton object was increased by switching to the “Transparent 0” color mode to allow better visibility of intracellular bacteria and nuclei. Bacterial DNA signal was identified based on colocalization with the GFP signal and rendered in cyan to distinguish it from host nuclear DNA. Images depicted in Fig. 4 were made with the snapshot function of Imaris at 600 dpi.

### Analysis of phagocytosis by RAW264.7 macrophages

*A. fumigatus* conidia were harvested from AMM agar plates, labeled with 0.1 mg/ml Fluorescein-isothiocyanate (FITC, Sigma-Aldrich, Germany) in 0.1 M Na_2_CO_3_ for 30 min and washed three times with PBS. *S. xylosus* MAM1901 and *M. sciuri* MAM2035 were harvested from TSA plates, and labeled with 4 µg / ml for *S. xylosus* and 10 µg / ml for *M. sciuri*, respectively, FITC in 0.1 M Na_2_CO_3_ for 5 min and washed three times with PBS.

RAW264.7 cells (ATCC:TIB-71) were cultivated in DMEM at 37 °C with 5% (v/v) CO_2_ in a humidified chamber. For infection experiments, RAW264.7 cells were seeded in 12 well plates at a density of 5 × 10^5^ cells per well. *L. murinus enop-GFP*, FITC-labeled *A. fumigatus* conidia, FITC-labeled *S. xylosus*, and FITC-labeled *M. sciuri* were added at an MOI of 0.5. Infection was synchronized by centrifugation for 5 min at 100 x g for *A. fumigatus*-infected cells and for 7 min at 600 x g for cells infected with bacteria. After 30 min incubation at 37 °C, 5% (v/v) CO_2_, cells were washed three times with PBS and detached using 0.5 mL of TrypLE (Thermo Fisher, Germany) per well for 5 min.

For fixation, the cell suspensions were centrifuged for 2 min at 600 g and 4 °C, resuspended in 4% (v/v) formaldehyde in PBS for 15 min at RT, and washed twice. Cell suspensions were stained with fresh concanavalin A AF647 (Invitrogen, USA) with a final concentration of 10 µg/ml. Imaging flow cytometry was conducted with Cytek Amnis ImageStreamX MkII (Cytek Biosciences, USA) using Inspire Software (Cytek Biosciences, USA), 5000-10000 cells were measured. Internalization analysis was performed with Ideas software (Cytek Biosciences, USA) using an Internalization Wizard. The gating strategy is described in detail in Fig. S3.

### Experimental schematics

Schematics of the experimental design were created using BioRender (https://biorender.com/).

### Quantification and statistical analysis

For the statistical evaluation of *L. murinus enop-GFP* abundance in lung samples determined by CFU count, the statistical analysis was performed using the unpaired t test with Welch’s correction. The Pearson correlation was used to determine correlation between CFU numbers and DNA abundance in the lung samples from days 0 and 1 post-colonization. Kruskal-Wallis test was used to statistically evaluate differences on internalization rate for RAW264.7 lung macrophages confronted with *L. murinus enop-GFP*, *S. xylosus*, *M. sciuri*, and *A. fumigatus*.

## Results

### A transgenic *L. murinus enop-GFP* strain was generated based on whole genome sequencing

To analyze whether members of the lung microbiome can persist in the lungs and form a resident community or represent a transient bacterial community tightly controlled by the host, we aimed to fluorescently label bacteria native to the lungs and analyze their lung colonization dynamics (Fig. 1A). *L. murinus* has been shown to be a common and dominating member of the murine lung microbiome and to possess immunomodulatory activity in the murine lungs [22, 23, 32–34]. Therefore, we used *L. murinus* MAM1902, a strain that we had previously isolated from the murine lower airways [32] as a representative lung commensal for our experiments. First, we carried out whole genome sequencing of *L. murinus* MAM1902. Overall, the genome size and GC content of the strain resembled that of the deposited genomes of the species. *L. murinus* MAM1902 did not contain plasmids. Sequencing and assembly of the circular genome of *L. murinus* MAM1902 resulted in a single, circularized 2,225,271-bp chromosomal genomic sequence with a GC content of 40.0 % (NCBI BioProject ID: 1433200). Genome annotation with the NCBI Prokaryotic Annotation Pipeline resulted in identification of 2,120 protein-encoding genes, 21 pseudogenes, and 86 genes for structural RNAs. Phylogenetic analysis using the whole-genome sequence mirrored the taxonomic assignment of the *L. murinus* MAM1902 strain at the species level (Fig. S4).

**Fig. 1.**
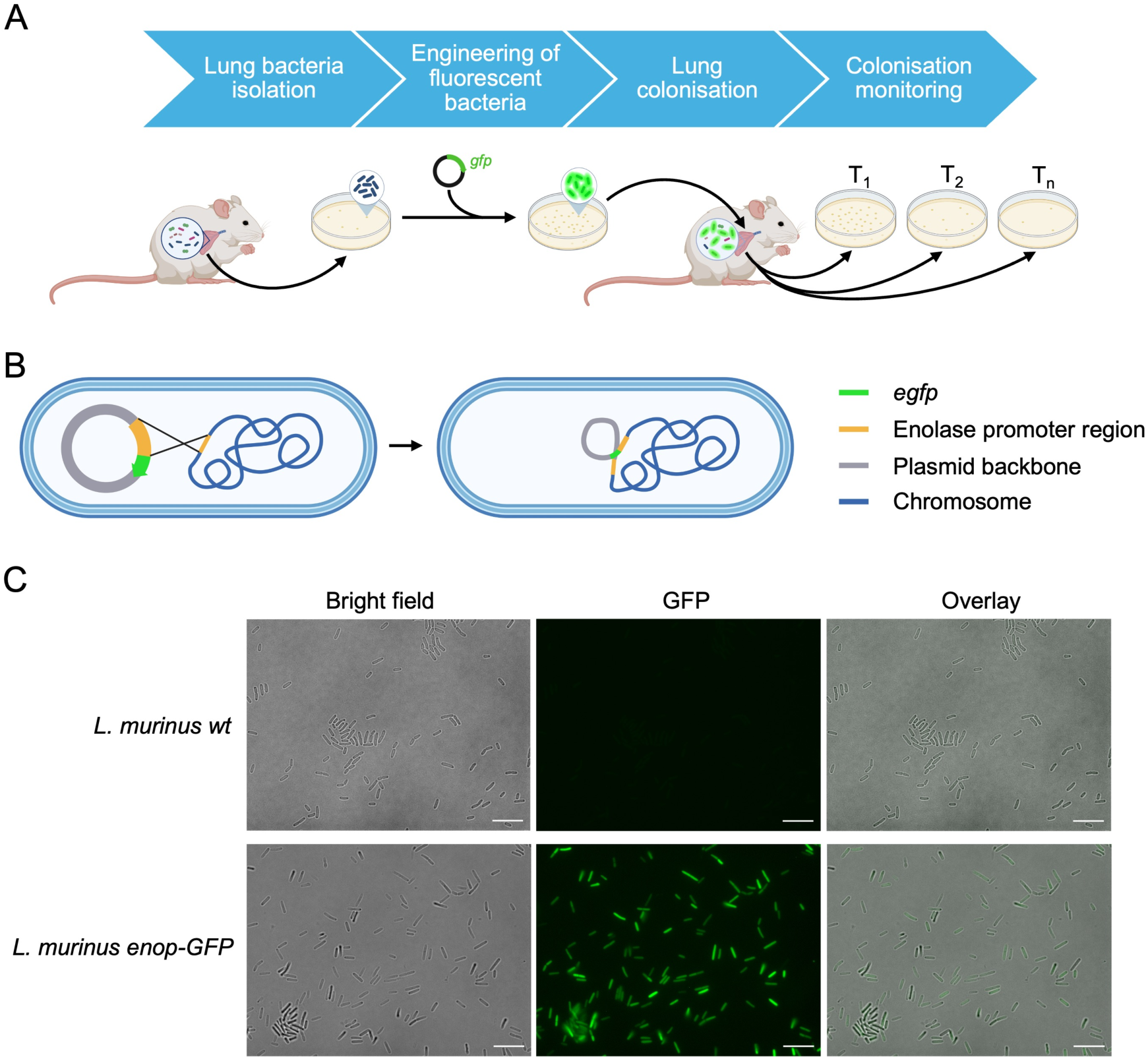
*L. murinus* was fluorescently labelled for monitoring presence of commensal bacteria in the lung. A) Experimental overview representing isolation of commensal bacteria from the murine lower airways, fluorescence labeling of the isolated bacteria by genetic engineering, administration of these bacteria to murine lungs, and monitoring the presence of bacteria at different time points over time. B) Schematic representation of the production of the GFP-producing strain *L. murinus enop-GFP*. In this strain, the *GFP*-encoding plasmid was inserted into the genomic enolase promoter region. C) Light microscopic images of *L. murinus* wild type and GFP-producing *L. murinus enop-GFP* strain cultivated anaerobically on CBA. Scale bar: 10 µm.

To investigate the lung colonization dynamics, we developed a strain that produced GFP, enabling us to track the bacteria over a longer period of time. For this purpose, GFP was initially expressed from a plasmid (pTRKH3-ermGFP; addgene plasmid # 27169 [41]). However, in the absence of erythromycin selection *L. murinus* lost the plasmid. Therefore, we generated a strain with a GFP gene integrated into the genome. For this, *L. murinus* was transformed with the vector pTRKH3-L.m.int.enoGFP which contained an integration cassette. The cassette contained the *GFP* gene under control of 1,000 bp enolase promoter region from *L. murinus* MAM1902 (1,467,752 bp to 1,468,751 bp), the chloramphenicol acetyltransferase gene as described by Vezina *et al.* [49], and was flanked by two homology regions (1,215,140 bp to 1,216,436 bp and 1,216,482 bp to 1,217,726 bp) (Fig. S5). Genome sequencing confirmed the genomic integration of the plasmid at the enolase promoter region (Fig. 1B). *L. murinus enop-GFP* stably produced GFP for several generations on Columbia Blood Agar (CBA). Furthermore, GFP expression (Fig. 1C) did not result in a fitness defect in MRS broth (Fig. S6).

### *L. murinus enop-GFP* is rapidly cleared from the mouse lungs

To track presence of commensal bacteria in the lungs overtime, firstly, 10^5^, 10^6^, 10^7^, and 10^8^ cells of *L. murinus enop-GFP* were administered intranasally to healthy mice and the presence of *L. murinus enop-GFP* in the lungs was analyzed two days after colonization by plating of dissociated lung samples on CBA (Fig. 2A). *L. murinus enop-GFP* was detected in the lungs of mice colonized with 10^7^, but not 10^6^ and 10^5^ bacterial cells, demonstrating that lower bacterial doses are cleared by the host within this short time period (Fig. 2B). Mice colonized with the highest dose exhibited severe lethargy and were sacrificed before the end of the experiment for animal welfare reasons. These results suggest that even non-pathogenic commensal bacteria may become harmful to the host when present above a certain abundance threshold.

**Fig. 2.**
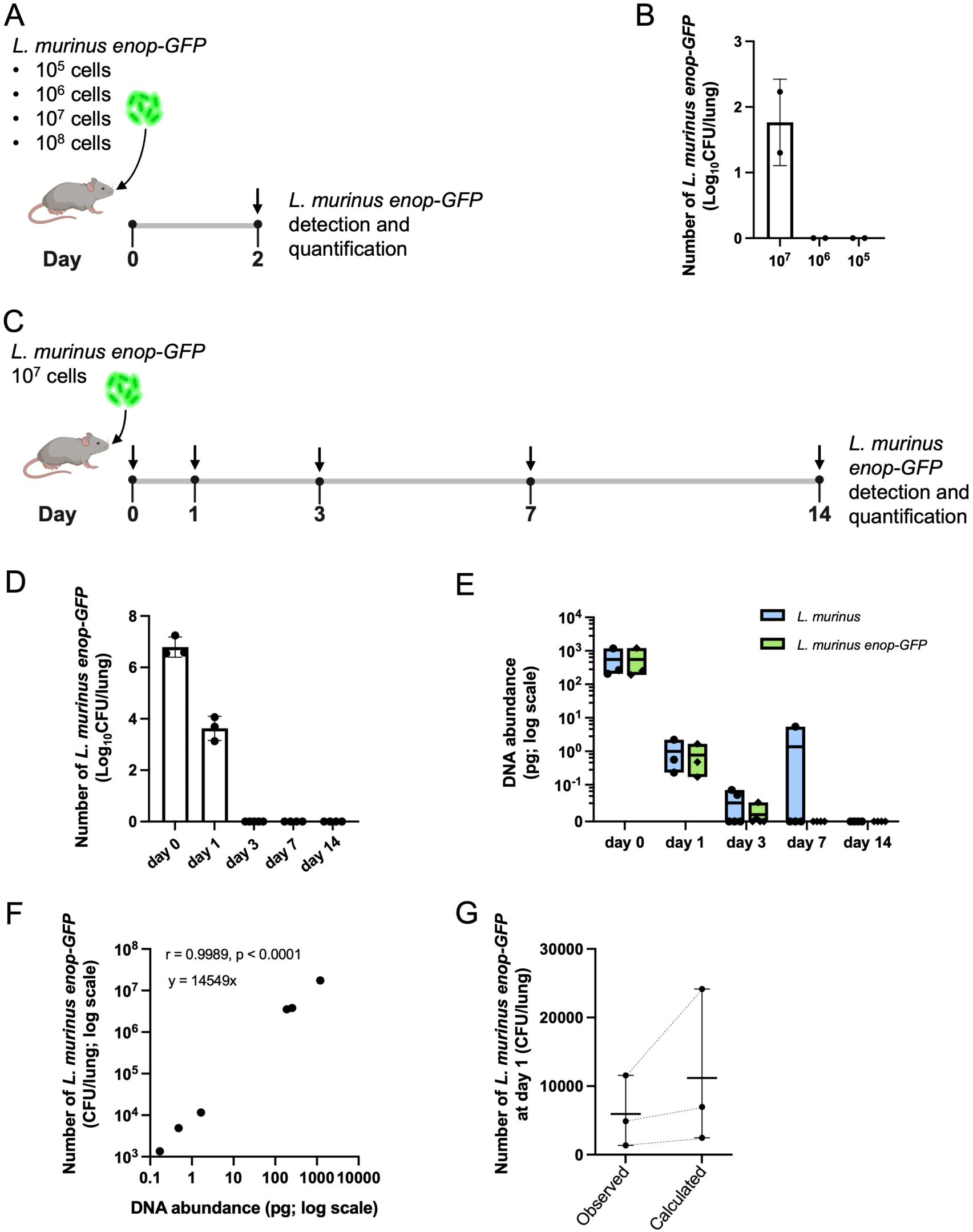
*L. murinus enop-GFP* is rapidly cleared from the mouse lungs. A) Experimental design for determination of the administration dose. *L. murinus enop-GFP* presence in the lungs was analyzed two days after administration of four different doses. B) *L. murinus enop-GFP* quantification in the lungs by CFU count; n = 2 (of note, CFU numbers were not determined in animals infected with 10^8^ bacteria as they had to be sacrificed before the end of the experiment). (legend continues on next page) C) Experimental design for the colonization analysis over 14 days. 10^7^ bacteria were administered intranasally to mice and *L. murinus enop-GFP* presence in the lungs was analyzed 5 minutes (day 0), 24 hours (day 1), 72 hours (day 3), 7 days and 14 days after administration. D) *L. murinus enop-GFP* quantification in the lungs at days 0, 1, 3, 7, and 14 by CFU count. E) *L. murinus* and *L. murinus enop-GFP* quantification in the lungs at days 0, 1, 3, 7, and 14 by qPCR. DNA quantity in picograms per mg of mouse DNA is shown. F) Correlation of *L. murinus enop-GFP* DNA quantities and CFU numbers in the lungs at days 0 and 1. The Pearson correlation was used to determine the correlation. G) Observed CFU counts and CFU values calculated from bacterial DNA abundance for *L. murinus enop-GFP* at day 1 post-administration. Calculation was performed with the equation Y = 14,549 × X based on the correlation data (F), where X represents DNA abundance (pg per 1 µg of mouse DNA) and Y represents CFU per lung. Mean and range are shown. D)– G) n = 3 for day 0 and day 1, n = 5 for day 3, and n = 4 for days 7 and 14.

To investigate whether *L. murinus enop-GFP* is able to persist in mouse lungs over a longer period of time, 10^7^ *L. murinus enop-GFP* cells were intranasally administered to mice and their presence in the lungs was analyzed 5 minutes (day 0), 24 hours (day 1), 72 hours (day 3), 7 days and 14 days after administration (Fig. 2C). For quantification of *L. murinus enop-GFP*, viable bacteria were enumerated by plating dissociated lung samples on CBA. Then, CFUs were counted. Additionally, *L. murinus enop-GFP* DNA was quantified by qPCR. From the samples obtained on the day of administration, on average, 8.257 x 10^6^ ± 7.966 x 10^6^ *L. murinus enop-GFP* CFUs were recovered. The CFU number dropped over 1,000-fold at day 1 post-colonization compared to day 0 (*p* = 0.001, unpaired t test with Welch’s correction), and no *L. murinus enop-GFP* colonies were detected at day 3 post-administration (Fig. 2D). Of note, only *L. murinus enop-GFP* was isolated from the lung samples on days 0 and 1. In contrast, *L. murinus* wild type and other bacterial colonies grew on plates inoculated with lung samples from days 3, 7, and 14 post-administration indicating bacterial immigration to the lungs.

To quantify *L. murinus enop-GFP* DNA as well as its proportion of the total lung *L. murinus* DNA, we performed qPCR on the gene encoding the *L. murinus* penicillin-binding protein and on the GFP-gene. The DNA quantification was performed in relation to the mouse DNA quantified by qPCR on the mouse beta-actin gene. For the samples from day 0 and day 1, *L. murinus enop-GFP* DNA abundance was almost equal to *L. murinus* DNA abundance (Fig. 2E). On day 3 post-administration, DNA of both wild-type *L. murinus* and *L. murinus enop-GFP* was detected at very low abundance in one sample, whereas in another sample only *L. murinus* DNA was detected (Fig. 2E). In both samples, wild-type *L. murinus* was isolated by plating. On day 7 post-administration, *L. murinus* DNA was detected in a single sample, while neither wild-type *L. murinus* nor *L. murinus enop-GFP* DNA were detected in lung samples collected 14 days post-administration (Fig. 2E). The presence of *L. murinus* DNA in samples from days 3 and 7 post-administration correlated with high numbers of bacterial colonies on agar plates, including colonies with *L. murinus* morphology. Consistently, when no or only few colonies with *L. murinus* morphology were observed, its DNA was not detected.

To estimate, if all detected *L. murinus enop-GFP* DNA indicated presence of viable bacteria, we performed a correlation analysis between CFU numbers and DNA abundance in the samples from days 0 and 1. As a result, a very strong correlation was observed (r = 0.9989, *p* < 0.0001, Pearson correlation) (Fig. 2F). The relationship was described by the linear function Y = 14,549 × X, where X represents DNA abundance (pg per µg of mouse DNA) and Y represents CFU per lung. Notably, this function was primarily dependent on the higher-value subset of samples obtained on day 0. When the equation was applied to DNA abundances from day 1, the predicted CFU numbers were 1.89-fold higher than the CFU counts recovered by culture, most likely reflecting the detection of DNA from both viable and nonviable bacteria rather than CFUs alone (Fig. 2G).

Taken together, for the highest well-tolerated dose of administered *L. murinus enop-GFP* we observed a strong decrease in numbers already 24 hours after administration and the number of bacteria was below the detection limit from day 3 post-administration. Our results indicate that 24 hours after administration around 50 % of the detected DNA belonged to dead *L. murinus enop-GFP* suggesting killing of the bacteria in the lungs.

### *L. murinus enop-GFP* cells are rapidly phagocytosed by lung immune cells

To analyze *L. murinus enop-GFP* interaction with host cells during lung colonization, microscopy and imaging flow cytometry analyses of lung samples of mice inoculated with 10^7^ *L. murinus enop-GFP* were performed. Microscopy of precision-cut lung slices (PCLS) revealed aggregates of individual *L. murinus enop-GFP* in alveolar spaces in all analyzed lungs on the day of administration and at day 1 post-administration (Fig. 3 and 4).

**Fig. 3.**
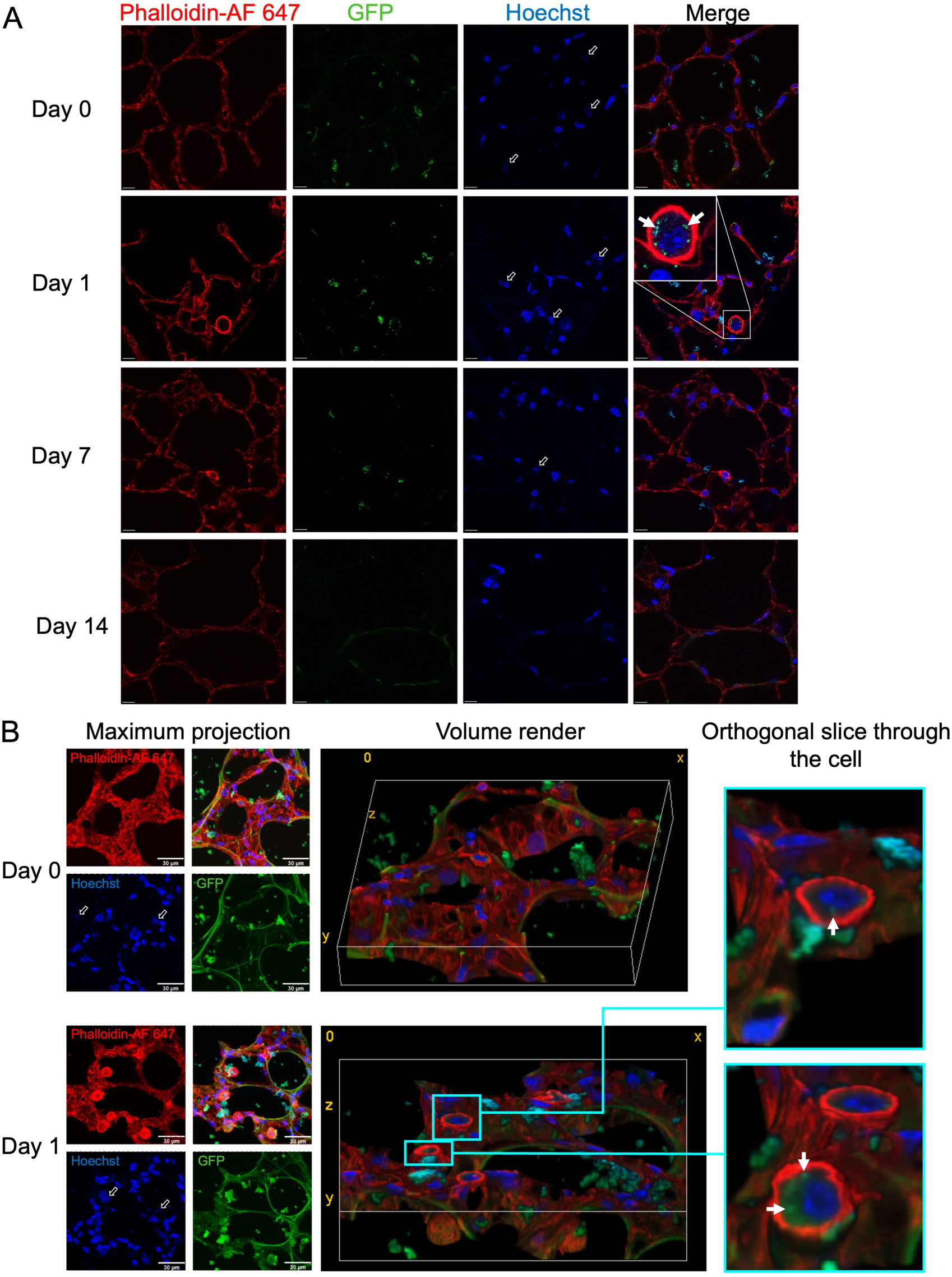
*L. murinus enop-GFP* is rapidly internalized by phagocytic cells, as detected in precision-cut lung slices. A) Confocal microscopy images of PCLS of mouse lungs inoculated with *L. murinus enop-GFP* analyzed at days 0, 1, 7, and 14 post-administration. Scale bar 10 µm. (legend continues on next page) B) Z-stacks visualized either as maximum projection, as a volume render with a bounding box size of 135×135×21.3 µm or as an orthogonal slice through the volume render (scaling not given due to zoom and rotation) to show internalization. Scale bar: 30 µm. For (A) and (B), bacteria were distinguished from autofluorescent cell matrix based on morphology and the overlap of DNA (Hoechst) and GFP signals. Hollow arrows in the Hoechst panel indicate exemplary clumps of Hoechst-stained bacteria. Putative phagocytic cells were identified based on size, internalization of bacteria and intensity of the cytoskeleton signal. Internalized bacteria are indicated by white arrows.

Rare bacterial aggregates were detected in the single analyzed lung at day 7 post-administration likely representing non-viable cells rather than persistent cells. Their presence may reflect incomplete early clearance, potentially associated with initially higher bacterial loads, which is in line with the substantial variability observed on the day of inoculation. No *L. murinus enop-GFP* signal was observed at the latest time point, which is consistent with the quantification results (Fig. 3A).

We did not observe any attachment or biofilm formation by *L. murinus eno-GFP* in the lungs. At day 1 post-administration, accumulation of cells with phagocytic morphology and very intense cytoskeleton signal were visible, many of which exhibited a signal from *L. murinus eno-GFP* in the cytoplasm (Fig. 3 and 4). This finding suggests recruitment of phagocytes to the place where bacteria were present, and their phagocytosis.

**Fig. 4.**
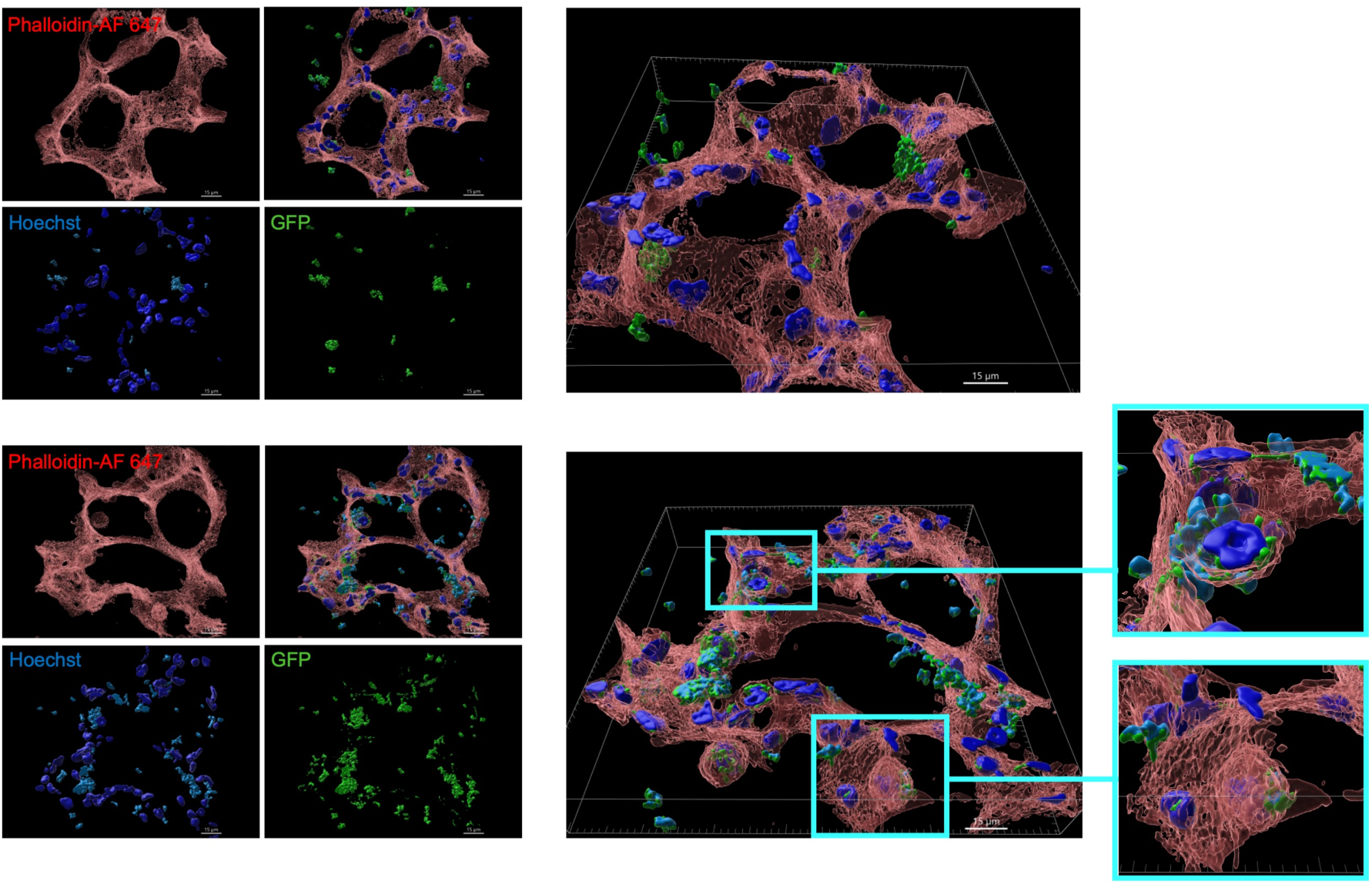
3D reconstruction of PCLS confirms internalization of *L. murinus enop-GFP* by phagocytic cells. Z-stacks were analyzed using Imaris software. Images were generated using the snapshot function of Imaris at 600 dpi. Scale bar: 15 µm. Bacteria were identified based on the overlap of Hoechst (cyan, bacterial DNA) and GFP (green) signals. Red: cytoskeleton (phalloidin-Alexa Fluor 647); blue: host DNA (Hoechst).

Analysis of internalization across the total lung cell population by imaging flow cytometry revealed a trend toward an increased number of cells with internalized bacteria at day 1 post-administration (Fig. 5A,B). This was consistent with our observations of an accumulation of cells exhibiting phagocyte-like morphology in microscopic pictures of PCLS.

**Fig. 5.**
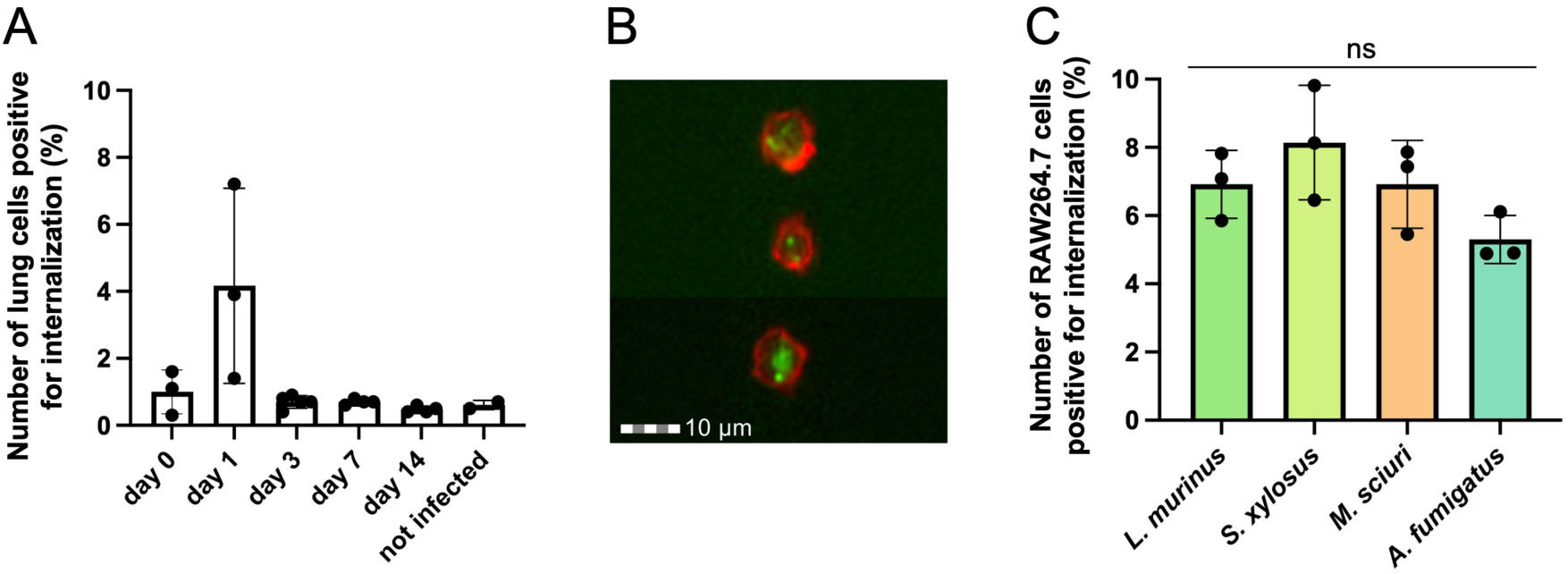
Flow cytometry reveals lung cell internalization of *L. murinus enop-GFP* within 24 h and macrophage phagocytosis. A) Quantification of *L. murinus enop-GFP* internalization by lung cells at days 0, 1, 3, 7, and 14 post-administration. Lungs were dissociated and the cell suspension was analyzed for internalization-positive events with imaging flow cytometry. B) Examples of internalization-positive cells. Red – cell membrane, green – GFP. C) Comparison of the internalization rate for RAW264.7 lung macrophages confronted with *L. murinus enop-GFP*, *S. xylosus*, *M. sciuri*, and *A. fumigatus* after 30 min with an MOI of 0.5. Difference was estimated using Kruskal-Wallis test.

To check if epithelial cells contributed to phagocytosis of *L. murinus enop-GFP* cells along with professional phagocytes as it has been shown for many lung pathogens [50], we quantified internalization by EpCAM-positive lung cells. We did not observe internalization of *L. murinus enop-GFP* by lung epithelial cells at any time point (Fig. S7).

### Irrespective of differences in their abundance in the lung, lung-derived bacteria are phagocytosed with the same rate by murine RAW264.7 macrophages

We observed that *L. murinus enop-GFP* was rapidly cleared from the lungs of mice, at high administration doses, likely due to phagocytosis by professional phagocytes. To address the question whether *L. murinus* is phagocytosed less rapidly than microorganisms found in lower abundances in the respiratory tract, we analyzed the phagocytosis rate by murine RAW264.7 cells of *L. murinus*, *Staphylococcus xylosus*, and *Mammaliicoccus sciuri*, as well as conidia of the lung-pathogenic fungus *Aspergillus fumigatus. S. xylosus* and *M. sciuri* were chosen as members of the murine lower airways microbiome found at different abundances – *S. xylosus* was primarily localized to the trachea with minimal presence in the lungs, whereas *M. sciuri* exhibited low abundance across both lower airway sites [32]. Because *A. fumigatus* conidia are widely distributed and readily inhaled, the fungus has been detected as a member of the lung mycobiome of healthy individuals [51–53]. Furthermore, *A. fumigatus*, as an opportunistic pathogen, is known to interfere with phagocytosis [54]. As a result, no difference was observed in numbers of internalization-positive cells between RAW264.7 macrophages confronted for 30 min at a multiplicity of infection (MOI) of 0.5 with *L. murinus enop-GFP*, *S. xylosus* MAM1901, *M. sciuri* MAM2035, and *A. fumigatus* (Fig. 5C).

Overall, our results indicate that *L. murinus* is rapidly cleared from the mouse lungs after administration, and that this clearance is driven at least in part by phagocytosis. In our experimental approach we did not observe major differences in the phagocytosis of high- and low-abundant microorganisms in healthy lungs and even compared to a fungal pathogen.

## Discussion

The microbiota plays a crucial role in mammalian development, physiology, and homeostasis. It contributes to the production of vitamins and other essential metabolites and modulates the function of our brain and immune system [16, 55]. Understanding the ecological organization of the microbiome at specific body sites is therefore essential for elucidating its functional roles and its impact on the host homeostasis. Our work builds upon earlier studies reporting a high prevalence of *L. murinus* in mouse lungs [23, 32–34]. Our results demonstrate that the dominant member of the murine lung microbiome, *L. murinus*, is rapidly cleared from the lungs following its introduction and is replaced by newly acquired bacteria, likely from the oral cavity. This finding indicates a rapid turnover of the lung microbiome and suggests the absence of a stable resident microbial community in the lung. This conclusion was supported by the detection of *L. murinus enop-GFP* within lung phagocytes, indicating active phagocytosis and suggesting that phagocytic clearance is a crucial mechanism restricting microbial colonization of the lungs.

Advances in sequencing technologies allowed detection of DNA of various microorganisms and viruses [3, 16], despite their low abundance in the lungs. Further, it has been shown that bacteria detected by the presence of their DNA are largely culturable and can be isolated [29]. Despite the very low bacterial load in the lungs, bacteria could be also visualized by *in situ* hybridization [33, 56, 57]. All these methods, however, are only a snapshot, as they are restricted to the detection of bacteria at a specific time point and do not allow the analysis of bacterial persistence in the host over time. This limitation can be overcome by labeling of lung-derived bacteria, prior to their administration, that allows tracking of spatial and temporal distribution of the introduced bacteria in the host [58–60]. We therefore generated a GFP-producing strain of *L. murinus*, that we had previously isolated from the lower airways of mice and that represents a common member of the murine lung microbiome [22, 23, 32–34]. This strain allowed to distinguish the intranasally administered bacteria from the native commensal bacteria in SPF mice and to detect and quantify them over time. Moreover, fluorescence microscopy enabled analysis of the localization of GFP-labeled *L. murinus* within the lungs and their internalization by phagocytic lung cells.

Quantification of viable cells and of DNA from GFP-expressing *L. murinus* indicated rapid clearance of this non-pathogenic bacterium from the mouse lungs. No viable bacteria were recovered beyond day 1 post-administration, consistent with DNA measurements. Consistent with our previous results [32], colonies of wild-type *L. murinus* were detected at days 3, 7, and 14 post-administration. As *L. murinus* is a member of the murine oral microbiome [61], it is highly probable that the wild-type *L. murinus* observed at later time points originated from the oral cavity.

Overall, our findings support the “adapted island model” proposed by Dickson *et al.*, based on the high interindividual variability combined with a low microbial load and spatial variation observed in lung samples of healthy individuals [25, 26]. This model predicts that bacterial composition in healthy lungs depends mainly on the dynamics of the microbial immigration to the lungs and their elimination, and less on microbial growth [26]. Our data substantiate this model as they strongly suggest that the lung microbiome is a highly dynamic microbial community with a rapid turnover of bacteria that entered the lungs.

According to our findings, the rapid turnover in the bacterial lung communities is mediated, at least in part, by the function of the immune system. It has previously been shown that alveolar macrophages and polymorphonuclear neutrophils (PMNs) are essential for the clearance of pathogenic bacteria such as *Staphylococcus aureus*, *Proteus mirabilis*, and *Pseudomonas aeruginosa* that were phagocytosed shortly after infection [62–64]. Furthermore, bacteria detected inside phagocytes were no longer viable [62]. Here, by quantification of viable bacteria and their DNA, we observed that approximately 50% of the *L. murinus enop-GFP* detected 24 h after administration at the highest well-tolerated dose were non-viable. This may reflect bacteria that had been phagocytosed and killed intracellularly. This is supported by the observation of *L. murinus enop-GFP* cells phagocytosed in the lungs at this time point. Association of lung bacteria with host cells has also been shown previously in lung transplant patients [65]. In particular, alveolar macrophages have been shown to represent around 80% of all cells detected in the BAL of healthy individuals and lung transplant patients without acute rejection or pneumonia [66, 67]. Therefore, it is very likely that the cell-associated bacteria in the lungs of the lung transplant patients were mostly represented by bacteria phagocytosed by alveolar macrophages. As reported here, microscopic analysis revealed that *L. murinus enop-GFP* localized intracellularly within cells displaying the morphology of phagocytic cells, which may represent alveolar macrophages, PMNs, and monocyte-derived macrophages.

An interesting question concerned the presence of bacteria in different abundances in the lung. It was conceivable that there were differences in their interaction with cells of the innate immunity causing different abundance. However, when we compared the phagocytosis of *L. murinus* with other bacteria found in the lung such as *S. xylosus* predominantly isolated from the trachea and only infrequently detected in lung samples and *M. sciuri* underrepresented at both sites of the lower airways [32], there was no difference in phagocytosis rate between these bacteria. This was even the case when the phagocytosis rates were compared to that of conidia of the lung pathogenic fungus *A. fumigatus*. This efficient phagocytosis of commensal bacteria likely contributes to maintaining the low microbial burden in the lungs, which has been reasoned to be an essential factor supporting the primary physiological function of the lungs, *i.e.*, gas exchange [3].

This data substantiate that the lung microbiome represents a highly dynamic and transient microbial community, where the acquired microorganisms are rapidly recognized and cleared by the immune system and are substituted by newly acquired microorganisms.

## Limitations of the study

The results obtained for quantification of the viable *L. murinus eno-GFP*, its DNA, and the internalization analysis could be directly compared as these analyses were performed on the same samples. However, this was not the case for microscopic analysis that had to be performed on separate lung samples due to sample preparation requirements, not allowing direct comparison of microscopic analysis with CFU and DNA quantification. Another limitation could be the relatively high administration dose of bacteria that might have induced a stronger immune response in the lungs compared to natural conditions. However, even at a low administration dose, no *L. murinus eno-GFP* bacteria were detected after two days, suggesting that hyper activation of the immune response is not required for clearance.

### Conclusions

The lung microbiome, despite its low biomass, contributes to modulation of host pulmonary homeostasis in both health and disease. The results of this study support the concept of the lung microbiome as a transient and highly dynamic community. They further suggest that rapid clearance of acquired bacteria by lung phagocytes is essential for maintaining pulmonary homeostasis, as increased abundance of even commensal bacteria can negatively affect host health. These findings are important for a comprehensive understanding of host-microbiome interactions in the respiratory tract and provide a foundation for future studies exploring the potential role of the lung microbiome for protection against lung infections.

## Declarations

### Ethics approval and consent to participate

Approval for the animal experiments was obtained from the responsible federal/state authority and the ethics committee, which are based on the provisions of the German Animal Welfare Act (permit no HKI-24-003).

### Consent for publication

Not applicable

### Availability of data and material

All materials within the paper are available from the corresponding author upon reasonable request.

The genome sequencing data have been deposited at the NCBI GenBank (BioProject PRJNA1433200) and will be publicly available as of the date of publication.

### Authors’ contributions

L.N., T.H., and A.A.B. designed the study; L.N., K.V., M.S., E.U., and T.H. performed experiments; L.N., K.V., S.S., Z.C., M.T.F., G.P., and A.A.B. analyzed and interpreted data; I.D.J., M.T.F., and G.P. provided analytic tools; A.A.B. acquired funding; L.N., T.H., and A.A.B. wrote the manuscript; K.V., S.S., Z.C., I.D.J., G.P. revised the manuscript. All authors read and approved the manuscript.

## Acknowledgements

We thank Sigrun Kirste, Andrea Hartmann, Sylke Fricke, and Christina Täumer for excellent technical assistance. We thank Dr. Lukas Radosa for help in the project administration.

## Supplemental information

Additional file 1. Supplemental figures, and Supplemental Table (PDF).

