## Additional file 1. Supplemental figures, and Supplemental Table (PDF). for "The murine lung microbiome is dynamic and transient"

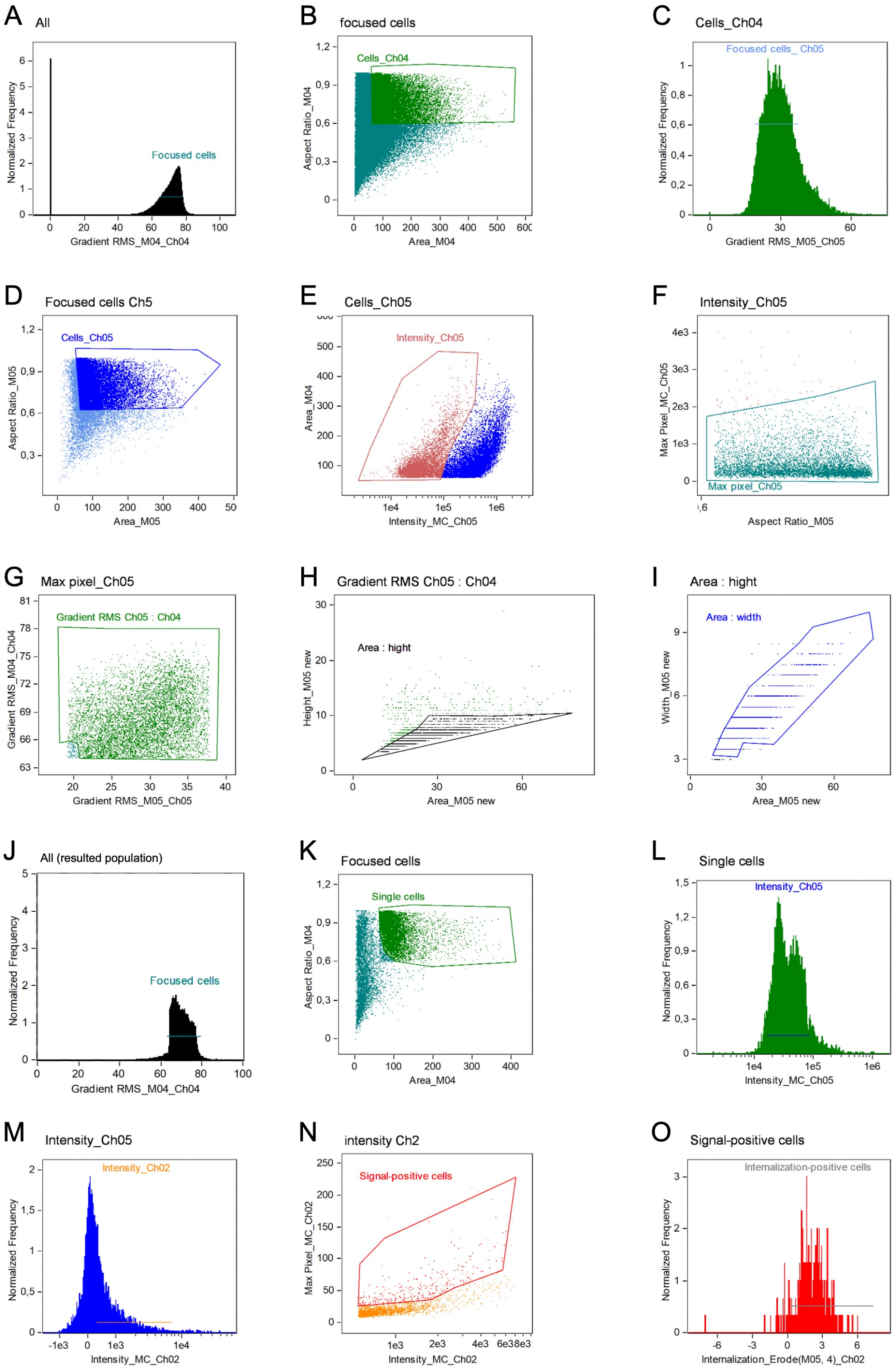


**Fig. S1. Gating strategy is applied to imaging flow cytometry data to quantify the lung cells stained with concanavalin A AF647 positive for *L. murinus enop-GFP* internalization.** (legend on next page)

(legend on next page)

**Fig. S1 (continued).** A) A population of focused cells gated based on high root mean square (RMS) gradient in the bright field channel. B) Single cells gated based on area and aspect ratio in the bright field channel. C) A population of focused cells gated based on RMS gradient in the channel 5 (642-745 nm). D) Single cells gated based on area and aspect ratio in the channel 5. E) Cell debris (higher signal intensity in the channel 5) removed based on area in the bright field channel and signal intensity in the channel 5. F) Cell debris (max pixel value in the channel 5) removed based on max pixel and aspect ratio in the channel 5. G) Cell debris removed based on its low RMS gradient values in both, bright field channel and channel 5. H) Cell debris removed based on area and height in the channel 5. I) Final cell population gated based on area and width in the channel 5. (J-O) Gating using an Internalization Wizard. J) From the obtained final population, a population of focused cells gated based on RMS gradient in the bright field channel. K) Single cells gated based on area and aspect ratio in the bright field channel. L) Single cells gated based on the intensity in the channel 5. M) Cells gated based on the signal in the channel 2 (505-560 nm). N) Cells with bacteria-like shaped signal gated based on the intensity and max pixel value in the channel 2. O) Internalization-positive cells gated based on the internalization erode.

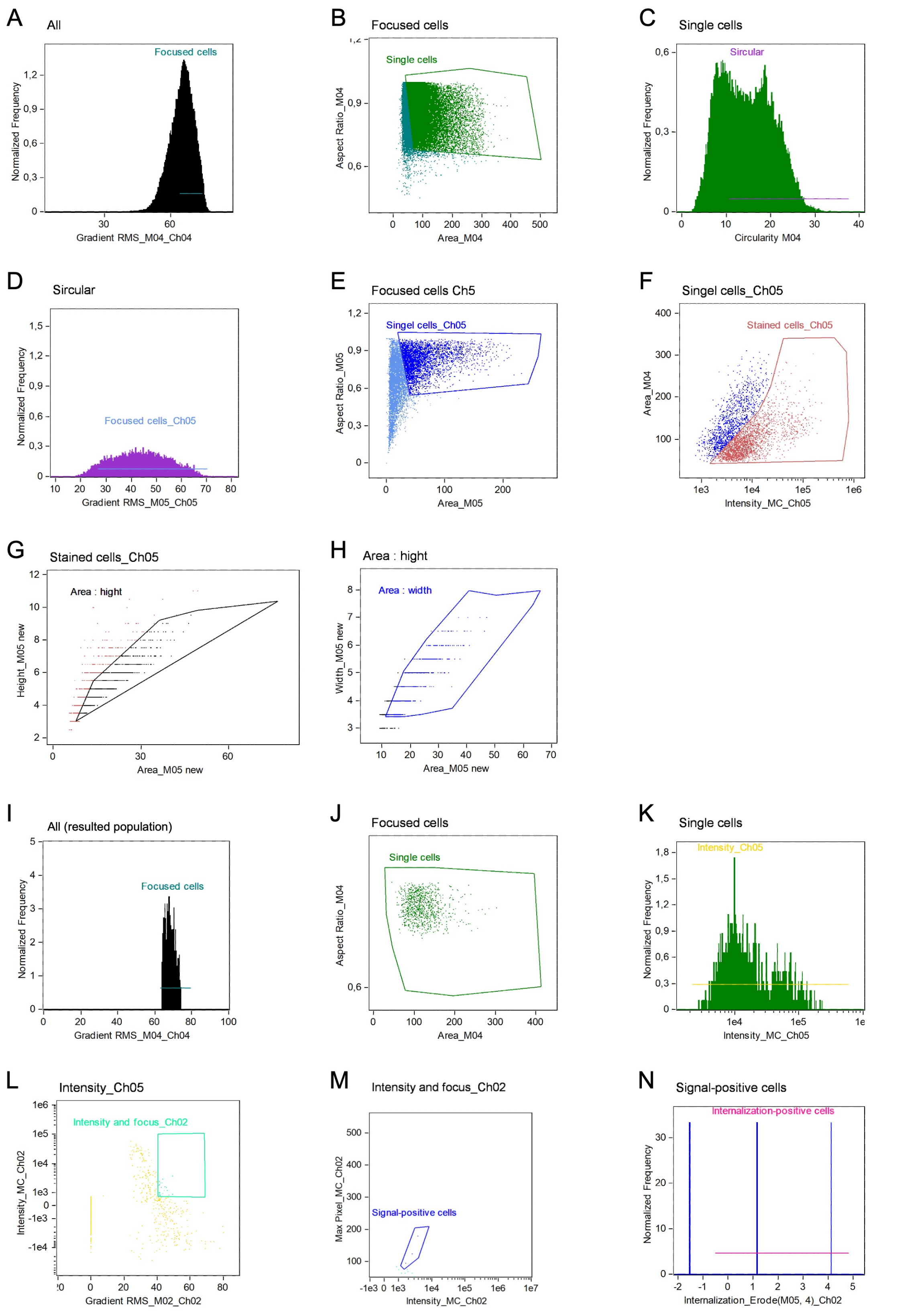


**Fig. S2. Gating strategy is applied to imaging flow cytometry data to quantify the CD326-positive lung cells positive for *L. murinus enop-GFP* internalization.** (legend on next page)

**Fig. S2 (continued).** A) A population of focused cells gated based on high root mean square (RMS) gradient in the bright field channel. B) Single cells gated based on area and aspect ratio in the bright field channel. C) Cells gated based on the circularity of the signal in the bright field channel. D) A population of focused cells gated based on RMS gradient in the channel 5 (642-745 nm). E) Single cells gated based on area and aspect ratio in the channel 5. F) Cells gated based on their area in the bright field channel and signal intensity in the channel 5. G) Cell debris removed based on area and height in the channel 5. H) Final cell population gated based on area and width in the channel 5. (I-N) Gating using an Internalization Wizard. I) From the obtained final population, a population of focused cells gated based on RMS gradient in the bright field channel. J) Single cells gated based on area and aspect ratio in the bright field channel. K) Single cells gated based on the intensity in the channel 5. L) Cells with bacteria-like shaped signal gated based on the RMS gradient and intensity in the channel 2 (505-560 nm). M) Cells with bacteria-like shaped signal gated based on the intensity and max pixel value in the channel 2. N) Internalization-positive cells gated based on the internalization erode.

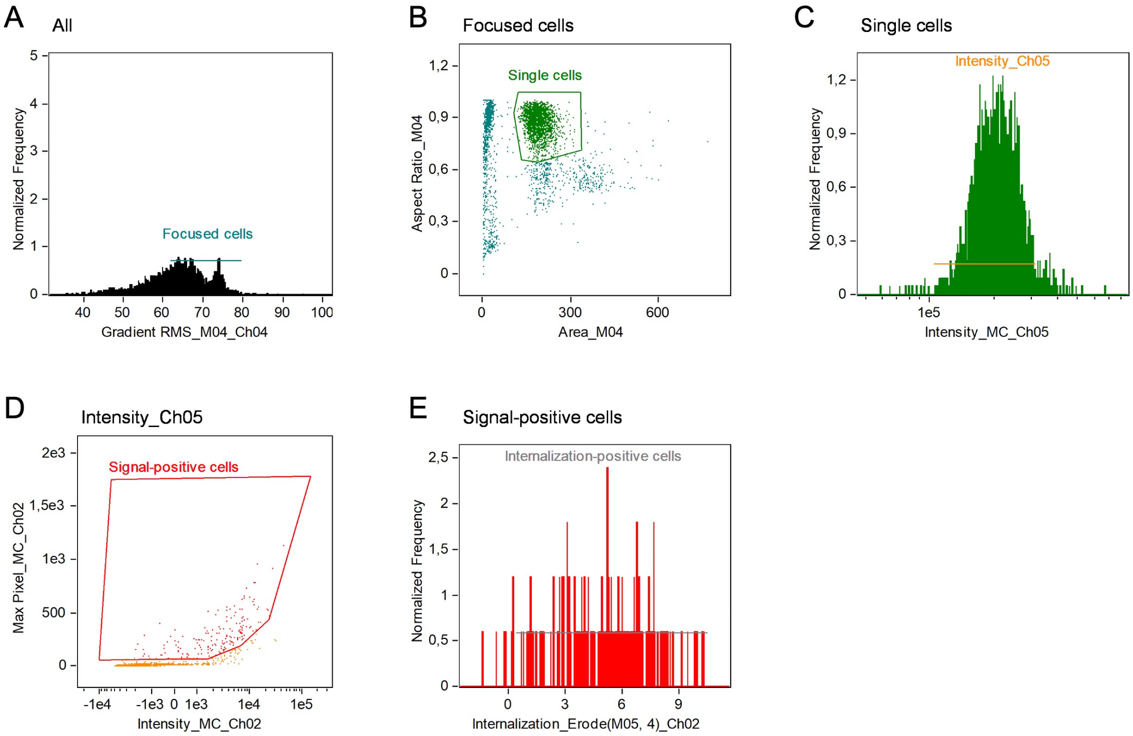


**Fig. S3. Gating strategy is applied to imaging flow cytometry data to quantify the RAW264.7 cells positive for internalization of fluorescently labeled microorganisms.**

(A-E) Gating using an Internalization Wizard. A) A population of focused cells gated based on RMS gradient in the bright field channel. B) Single cells gated based on area and aspect ratio in the bright field channel. C) Single cells gated based on the intensity in the channel 5 (642-745 nm). D) Cells with bacteria-like shaped signal gated based on the intensity and max pixel value in the channel 2 (505-560 nm). E) Internalization-positive cells gated based on the internalization erode.

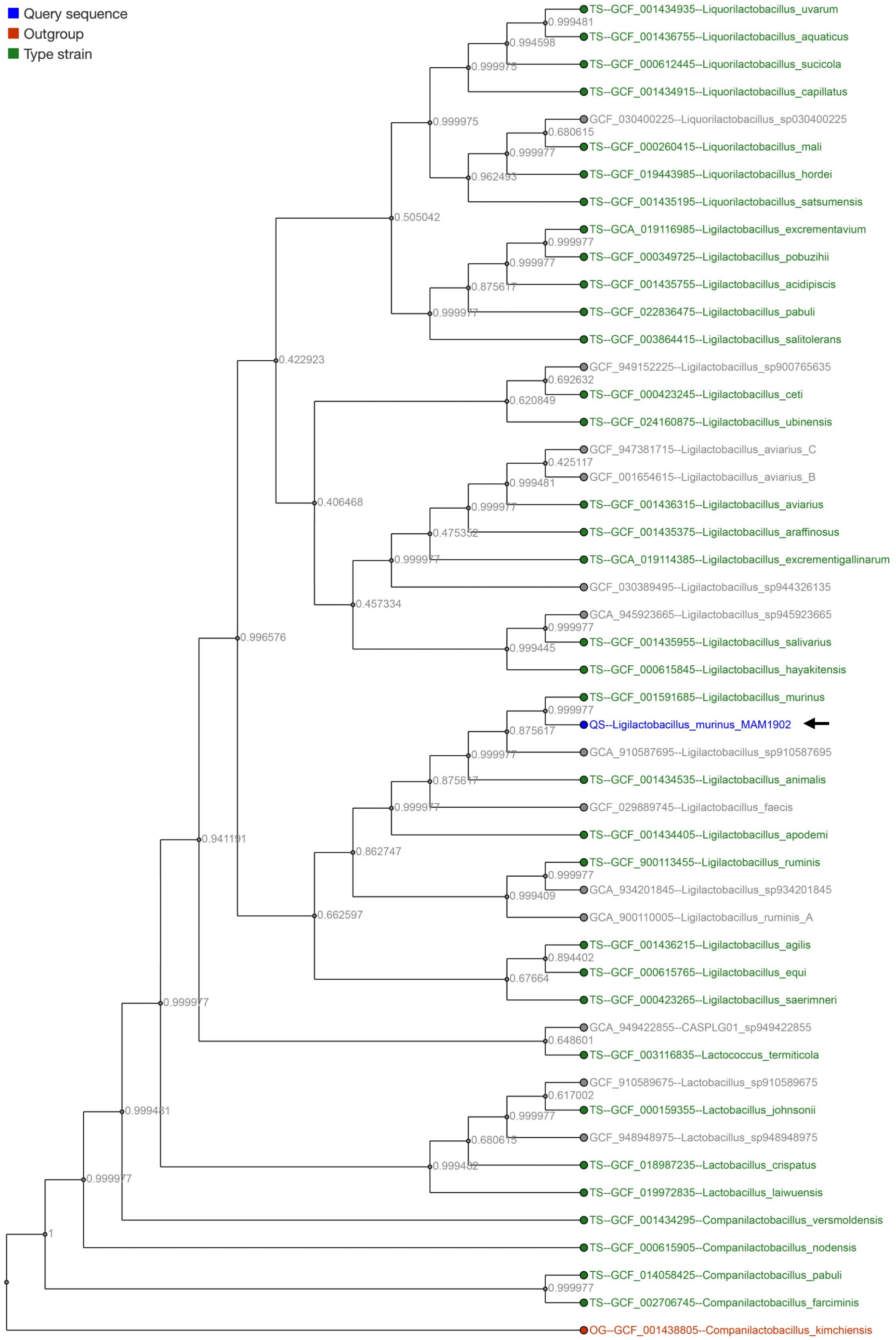


**Fig. S4. Phylogenetic analysis of *Ligilactobacillus murinus* MAM1902 based on the whole genome sequence mirrored the taxonomic assignment of the *L. murinus* MAM1902 strain at the species level.**

Analysis was done with AutoMLST2 in a Denovo mode and with standard settings.

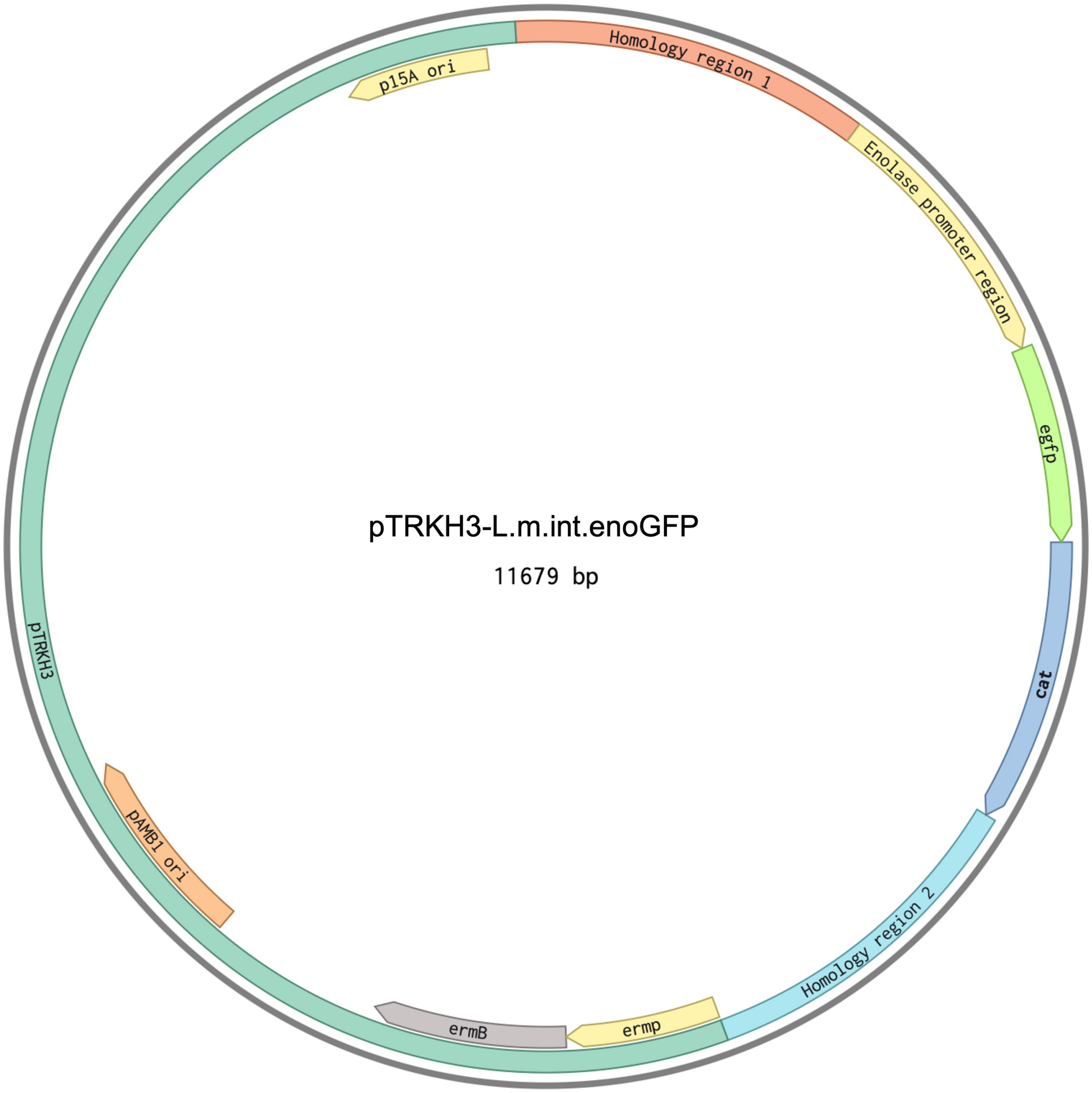


**Fig. S5. pTRKH3-L.m.int.enoGFP vector was used for GFP integration into *L. murinus* genome.**

Vector consists of a pTRKH3 backbone with erythromycin-resistance marker (*ermB*) and an integration cassette with two 1,200-1,300 bp long homology regions located in the inactive prophage region of *L. murinus* genome (1,215,140 bp to 1,216,436 bp and 1,216,482 bp to 1,217,726 bp), chloramphenicol acetyltransferase gene, and *egfp* under 1,000 bp-long enolase promoter region identified from *L. murinus* MAM1902 genome (1,467,752 bp to 1,468,751 bp).

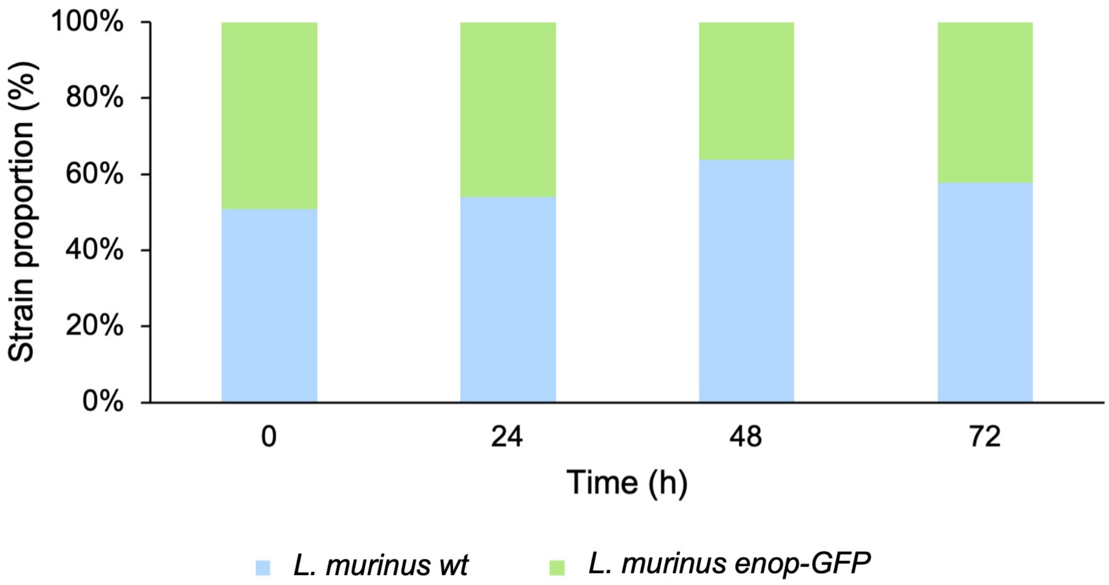


**Fig. S6. *L. murinus enop-GFP* fitness is comparable to that of the non-transformed wild-type strain.**

Fitness of *L. murinus enop-GFP* analyzed by its co-cultivation with *L. murinus wt* in MRS. *L. murinus enop-GFP* and *L. murinus wt* were quantified in the co-culture by CFU enumeration at the starting point, after 24 h, 48 h, and 72 h of cultivation. Total number of CFUs of *L. murinus enop-GFP* and *L. murinus wt* from one sample is set to 100%.

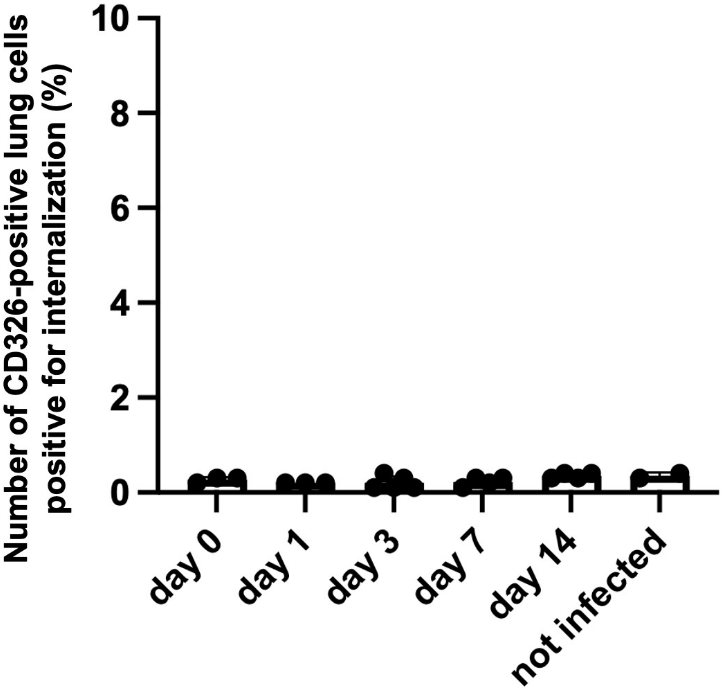


**Fig. S7.** ***L. murinus enop-GFP*** **is not internalized by the CD326-positive lung epithelial cells.** Quantification of *L. murinus enop-GFP* internalization by the CD326-positive lung cells at days 0, 1, 3, 7, and 14 post infection. Lungs were dissociated, the single cell suspension was incubated with CD326 (EpCAM) antibodies, and analyzed for internalization-positive events with imaging flow cytometry.

**Table S1. Oligonucleotides used in this study.** Bold letters mark homologous sequences for Gibson cloning.

| Name | Sequence and modifications |
|---|---|
| Lm_Peno_F | **CTCGTAAATCTTATTTGTTTAAGATTATC**TCTGACAAGTAATAAC |
| Lm_flank_2_R | GATCAGAACAGAAGTTAATTACTTTTACTCTAAAACTAAGC |
| Lm_flank_1_F | GAAATCAATGTTTGATTTTATTAGTTTTATACTCATTAAGACG |
| Lm_flank_1_R | GATAATCTTAAACAAATAAGATTTACGAGTAGCTATATTAAGG |
| pTRKH3_flank_1_F | **CCCATACGATATAAGTTGTAATTCTCATGT**CCAACAATAATAAAC |
| pTRKH3_flank_2_R | **GATGGC**TAGATGTAAGTGGTCGAGTC |
| Flank_2_pTRKH3_F | **TAAAATTTAAGACTCGACCACTTACATCTA**GCCATCTCCAGCAGC |
| pTRKH3_R | ACATGAGAATTACAACTTATATCGTATGGGGCTGACTTCAGGTGC |
| L.m.PBP_F | AAGCTTGGCGCATCGTCATC |
| L.m.PBP_R | CAACCGTGCCGTTCAAACTG |
| L.m.PBP_probe | 6-FAM/CAGGTGCAT/ZEN/AATCCATCAAAGGCTTAGCCGTAGAA/IBkFQ |
| EGFP_F | ATACAACTACAACTCCCACAAC |
| EGFP_R | TGGTAAAAGGACAGGGTCATC |
| EGFP_probe | 6-FAM/ TCAAAGCCA/ZEN/ACTTCAAGACCCGCCA/IBkFQ |
| b_actin_F | AGCACAGCTTCTTTGCAGCTCC |
| b_actin_R | TGGTGTCCGTTCTGAGTGATCC |
| b_actin_probe | 6-FAM/AGCGGGCCT/ZEN/TCGCTCTCTCGTGGCTAGTA/IBkFQ |
